# Fast remote nucleotide sequence alignment with Riboseek

**DOI:** 10.64898/2026.07.31.741718

**Authors:** Sukhwan Park, Kieran Didi, Andrew Favor, Anton Bushuiev, Soohyun Kim, Milot Mirdita, Martin Steinegger

## Abstract

Structure prediction has reached RNA, where generating deep alignments is now the bottleneck. We developed Riboseek, a search and alignment tool for RNA and DNA that represents sequences as overlapping di-mers. Riboseek is more sensitive for homology detection than BLASTN and nhmmer, and generates alignments 250- and 376-fold faster than nhmmer and rMSA. With structure-aware realignment, it approaches rMSA’s base-pair recovery, while remaining 29-fold faster. We provide a web server and 1.7 million precomputed alignments for putatively structured RNAs. Riboseek is freely available at https://github.com/steineggerlab/riboseek.

---

Nucleotide homology search plays an important role in genome annotation, comparative genomics, and RNA structure predictions. As public sequence repositories such as GenBank (1) and the European Nucleotide Archive (2) continue to grow, the practical value of a search method heavily depends on its speed vs. sensitivity tradeoff.

On one end of this tradeoff, fast tools such as BLASTN (3, 4) rely on heuristic word-hit seeding and extension, making their sensitivity dependent on seed length, extension parameters, and the presence of sufficiently conserved local matches (4, 5). As a result, distant nucleotide homologs can be missed when substitutions, indels, or sequence masking leave too few seedable regions.

On the other end, profile-based methods offer increased sensitivity by replacing a single query sequence with a statistical model of a sequence family. The current gold-standard tool nhmmer extended this framework to DNA/RNA homology search and improved detection of remote nucleotide homologs (6). Nevertheless, profile hidden Markov model (profile-HMM) nucleotide search is more than two orders of magnitude slower than the fastest seed-based approaches, which limits its scalability to large databases and many queries.

Another approach on the sensitive end relies on the fact that many RNA families conserve secondary structure through compensatory substitutions, allowing primary sequence to diverge while base-pairing interactions are maintained (7, 8). Covariance models capture this signal by jointly modeling sequence conservation and structural covariation in a single probabilistic framework (7), and their implementation in Infernal (9) has made them the standard for high-sensitivity RNA family detection in resources such as Rfam (10). Infernal’s filter pipeline and HMM-banded covariance-model alignment substantially improved the speed of covariance-model search, but the method remains highly computationally demanding. Consequently, Infernal is often the most sensitive choice when a trusted structural alignment is available, but it is less practical for a first-pass search of very large nucleotide datasets.

RNA MSAs have also become an input to structure prediction, where alignment depth affects model accuracy. AlphaFold 3 (11) and Protenix-v2 (12) both accept RNA MSAs, and the AlphaFold 3 pipeline builds them by searching RNAcentral (13) and NT (14) with nhmmer. An alternative and even slower approach is taken by rMSA (15), which combines nhmmer and BLASTN searches with Rfam-guided and structure-aware filtering. Because MSA generation based on large databases occurs for each query and can take hours, it becomes the main bottleneck in large-scale prediction.

Tools such as MMseqs2 (16) have transformed the speed– sensitivity tradeoff for protein homology search and enabled applications including protein structure prediction with AlphaFold2 (17) and ColabFold (18). No equivalent advance has been achieved for nucleotide sequences. Here, we present Riboseek, a sequence-based search engine that combines profile-based di-mer scoring, fast prefiltering and gapped alignment to identify homologous sequences at scale (Fig. 1). Across our benchmarks, Riboseek improves sensitivity over BLASTN, nhmmer and pairwise Smith–Waterman search while remaining two orders of magnitude faster than nhmmer and Infernal. Riboseek reads the input query as di-mers: two adjacent nucleotides define a single state. It then applies a di-mer substitution matrix (Methods) to construct a query profile, which is defined as its position-specific scoring matrix (PSSM). The profile extends each query di-mer position to related di-mer states (1a). Because nucleotide homologs may occur on either strand, Riboseek constructs forward and reverse-complement query profiles before searching the target database.

**Fig. 1.**
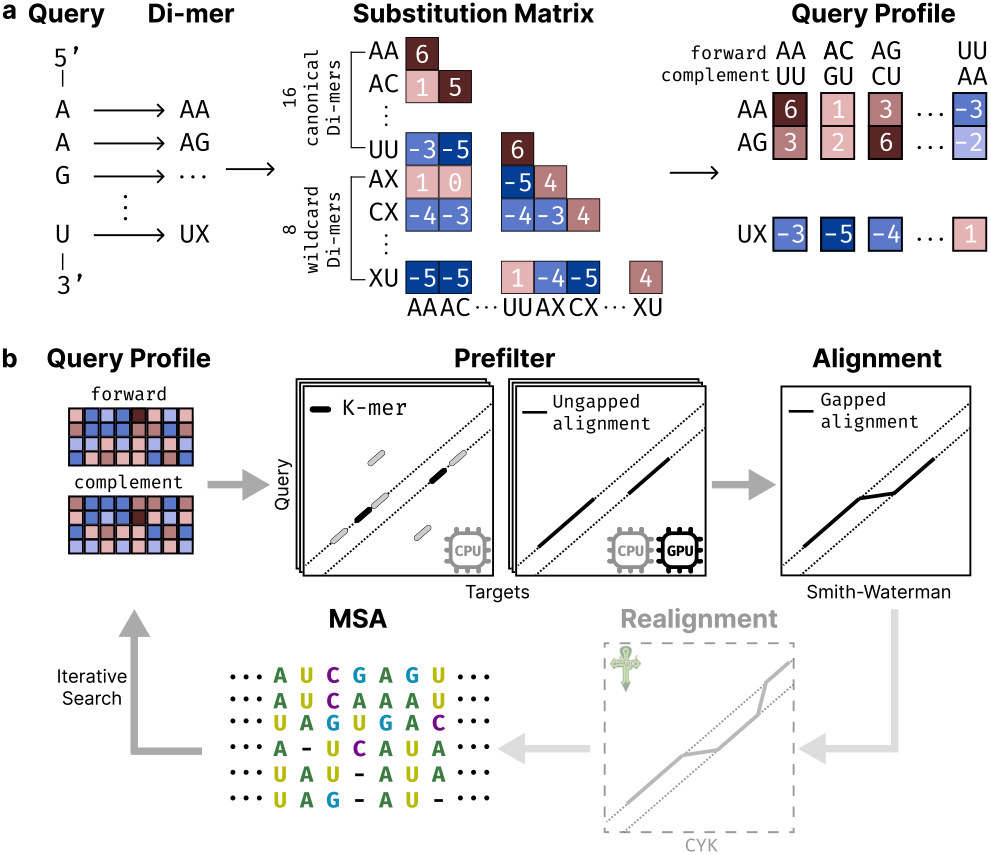
Riboseek workflow. **a**, Riboseek converts a nucleotide query into overlapping di-mers and uses a learned substitution matrix to construct forward and reverse-complement query profiles. The alphabet contains 16 canonical and eight wildcard di-mers. **b**, Candidate targets are identified using a *k* -mer-based or ungapped prefilter and aligned using affine-gapped Smith–Waterman-Gotoh alignment. In iterative searches, matched sequences are combined into a query-centered multiple sequence alignment (MSA) to update the query profile. Optionally, hits are realigned with Infernal’s CYK algorithm using a covariance model constructed from the Riboseek MSA and a predicted RNA secondary structure.

The search itself is organized as multiple steps (1b). In the first stage, Riboseek uses a fast *k*-mer or GPU-accelerated ungapped prefilter (19) to identify database entries that share short di-mer features with the query profile. Candidates that pass prefiltering are fully aligned using the affine gapped Smith–Waterman–Gotoh algorithm (20, 21).

Riboseek further improves sensitivity through iterative profile search. After an initial search round, matched target sequences are collected and used to build a multiple sequence alignment (MSA) and a position-specific scoring matrix (PSSM) query profile for the next search round. Riboseek also supports an optional structure-aware realignment with the Cocke–Younger–Kasami (CYK) algorithm using a covariance model built from the Riboseek MSA. For this purpose, we embedded Infernal as a library in Riboseek for the covariance model generation and structure-aware alignment.

To compute Riboseek’s substitution matrix, optimize its parameters and benchmark it, we first split Rfam 14.10 (10) families into training, validation and test sets, ensuring the sets do not share Rfam clans (Methods). The substitution matrix was then estimated from the training set of 3,235 families. For RNA benchmarking, we constructed a target database from the 155 test-set families in which each sequence has known homologs (same family) and non-homologs (different clan, or shuffled sequences), with random pseudogenomic flanks appended to every target to approximate genomic search conditions (Methods). As queries, we used the longest sequences from 71 test-set families with more than 20 members, and benchmarked the sensitivity and runtime of Riboseek against Infernal, BLASTN, nhmmer, and pairwise Smith– Waterman (SW). All tools received only the query sequence; Infernal’s model was built from it with an RNAfold-predicted secondary structure (22).

On this Rfam benchmark (Fig. 2a, b; Supp. Fig. 1), comparing tools at their first iteration, Infernal was the most sensitive, followed by Riboseek, which was more sensitive than nhmmer, SW, and BLASTN. To simulate realistic conditions of querying a large database, we timed searching the queries against RNAcentral v26 (Fig. 2b, inset). Riboseek’s first iteration completed in under two minutes, 544*×* and 777*×* faster than nhmmer and Infernal, respectively, 36.5*×* faster than nhmmer run as multiple parallel jobs, each using one CPU core, and only 2.9*×* slower than BLASTN. Without a GPU, Riboseek’s *k*-mer prefilter completed the same search in 30 minutes at the same sensitivity (mean ROC1-AUC 0.323), still 37*×* and 53*×* faster than nhmmer and Infernal. Two Riboseek iterations achieved the highest sensitivity of all methods, surpassing single-pass Infernal and the evaluated nhmmer iterations, while remaining 394*×* faster than Infernal.

**Fig. 2.**
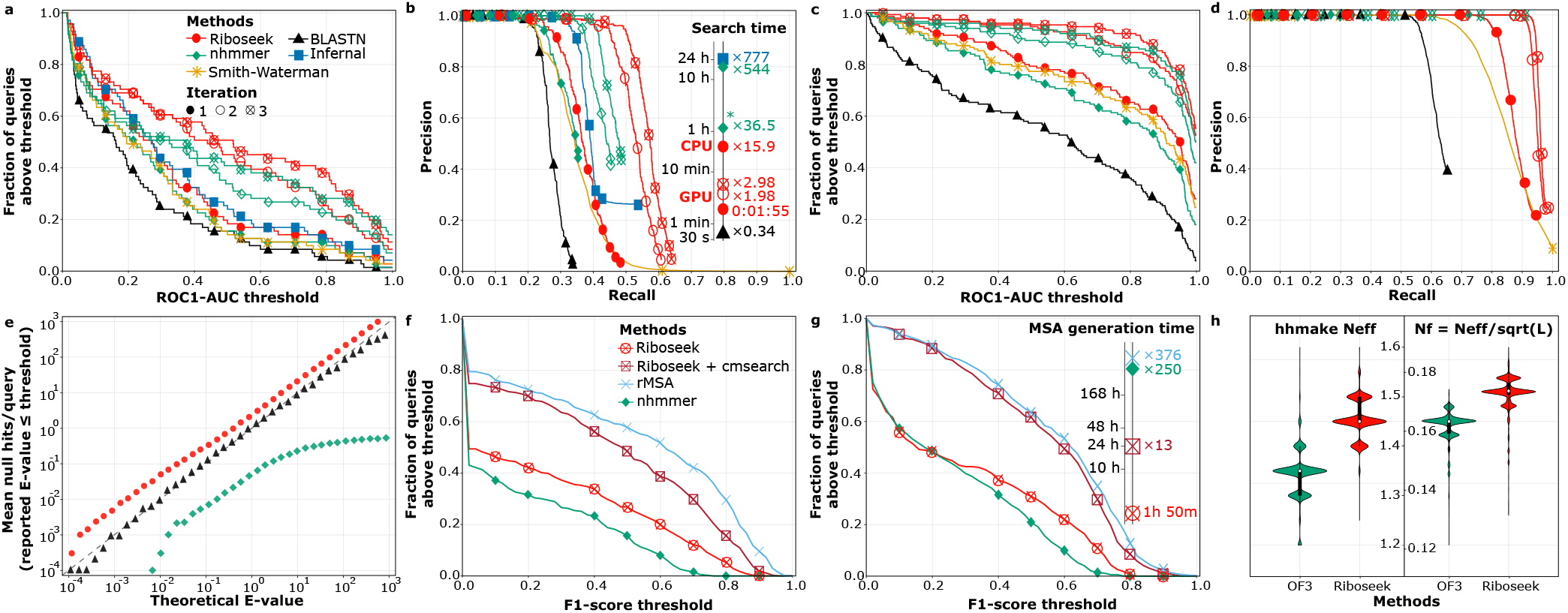
Benchmark results. **a, b**, Rfam benchmark: 71 test sequences were queried against 96,701 targets. For each query, true positives (TP; same-family hits) and false positives (FP; different-clan or shuffled sequence) were determined, ignoring same-clan hits. A query’s ROC1-AUC is the fraction of TP ranked above the first FP. **a**, The fraction of queries (y axis) above each ROC1-AUC threshold (x axis). **b**, Precision–recall curves for the same searches. The inset reports search runtimes for the same queries against RNAcentral v26 (Methods). Asterisk indicates the mean runtime of parallel jobs, with each job using one CPU thread. **c, d**, Dfam benchmark: 150 test sequences were each queried against a family-specific target database, treating family members as TPs and shuffled sequences as FPs. Panels as in **a, b. e**, E-values under the null: hits of shuffled RNAcentral sequences. For each theoretical threshold (x axis), the mean number of observed hits per query with a smaller reported E-value is shown (y axis). Dashed line (y = x) indicates the expected relationship if E-values are well calibrated. **f, g**, Secondary-structure benchmark: 346 test sequences with reference base pairs were queried against RNAcentral and NT using each tool’s pipeline. Resulting MSAs were passed to R-scape (**f**) or plmc (**g**) to predict base pairs and compare them to the reference. For each F1-score threshold (x axis), the fraction of queries above that threshold is shown (y axis). The inset reports the total MSA-generation time. **h**, MSA diversity of 4,991 queries shared between the OpenFold3 RNAcentral and Riboseek collections. Effective sequence counts were computed with hhmake; 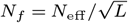, where *L* is the query length.

To attribute Riboseek’s sensitivity to individual components, we excluded the prefilter and repeated the benchmark with exhaustive Smith–Waterman–Gotoh alignment and optimized gap penalties. Di-mer scoring raised the mean ROC1-AUC over a mononucleotide alphabet from 0.286 to 0.298, matching the BlastR di-mer matrix (0.298), which was previously found to increase sensitivity of ncRNA search (23), and composition-bias correction raised it further to 0.331 (Supp. Fig. 2).

To benchmark DNA-family recognition, we produced a test set of 150 DNA families (Methods), derived from Dfam 2.0 (24). Each family representative was queried against its own database, where family members served as known homologs and shuffled sequences as non-homologs. Riboseek achieved higher sensitivity than BLASTN, nhmmer, and SW (2c, d).

Riboseek’s E-values are computed using the parameters of an Extreme Value Distribution fitted to the scores of null hits, with a post hoc correction (Methods). We evaluated the calibration of E-values by Riboseek and the other tools by comparing the distribution of reported E-values among null hits, generated from shuffled RNAcentral query sequences (Methods), to theoretical E-value thresholds. Riboseek and BLASTN were well calibrated, with reported E-values closely matching the number of null hits expected at each threshold (Fig. 2e). In contrast, nhmmer was conservative, reporting fewer null hits than expected per E-value cutoff.

We next benchmarked Riboseek against rMSA and nhmmer for their ability to generate MSAs for 346 RNA queries (Methods), which served as input for secondary structure prediction (Fig. 2f, g). Here, all tools searched RNAcentral v26 (30 gigabases) and NT (3.2 terabases). For a batch of 70 queries, formed by replicating seven sampled queries tenfold, Riboseek completed the search in 1.8 h, whereas nhmmer and rMSA required 250*×* and 376*×* longer, respectively (19.1 and 28.8 days); their runtimes were extrapolated from single-query runs, whereas Riboseek parallelises across queries and searched the batch in one run. Adding CYK realignment increased Riboseek’s runtime to 23.6 h, still 19.5*×* and 29.3*×* faster than nhmmer and rMSA, respectively (Fig. 2g, inset). At this target database size, runtime is dominated by the pre-filter, which Riboseek executes as a data-parallel kernel on a single NVIDIA RTX PRO 6000 GPU with 24 CPU cores; nhmmer and rMSA have no comparable GPU implementation and ran on 64 CPU cores.

Although rMSA achieved the highest secondary-structure prediction accuracy on this benchmark, Riboseek gained substantial accuracy after CYK-based realignment of its hits (Fig. 2f, g). This improvement suggests that Riboseek recovered a substantial number of true homologs, even when their initial MSAs required further refinement.

To assess Riboseek MSAs’ contribution to RNA tertiary structure prediction, we measured the mean local Distance Difference Test (lDDT) (25, 26) and the template modeling (TM) score (27) of PDB chains released after the training cutoffs of the Protenix-v2 and AlphaFold 3 models to their predictions. These chains were clustered and classified as structurally similar to the RNA chains used for model training (n=13) or as novel (n=15; Methods). With Protenix-v2, the better-performing of the two prediction models (Supp. Tables 1 and 2; Supp. Figs. 3–6), Riboseek MSAs improved over using nhmmer MSAs on the similar chains (mean lDDT 0.707→0.726, TM-score 0.551→0.593) and on the novel chains (equal lDDT of 0.614, TM-score 0.316→0.331). Both Riboseek and nhmmer improved substantially over single-sequence input.

To provide RNA MSAs at scale, we used Riboseek to pre-compute alignments for 1,731,677 RNA sequences predicted to form ordered secondary structures (Methods). We compared these with the RNAcentral and Rfam MSA collections released with OpenFold3 (28), encompassing 126,778 RNA sequences, 13.7-fold fewer than the Riboseek collection. For queries of similar median length (73 nt), Riboseek MSAs had a median depth of 64,379 sequences, versus 5,653 and 5,313 for the OpenFold3 RNAcentral and Rfam collections, respectively (11–12× deeper). For 4,991 queries shared between the Riboseek and OpenFold3 RNAcentral collections, Riboseek MSAs also had higher median *N*_eff_ and *N*_*f*_ values (Fig. 2h). Finally, we provide a Riboseek web API, integrated into ColabFold, which generates MSAs on demand for sequences not represented in the 1.7 million collection (Methods).

In conclusion, Riboseek broadens the accessible speed–sensitivity range for nucleotide search: it can rapidly identify candidate homologs in NT-scale resources with optional structure-aware refinement. Riboseek generates homolog sets and MSAs for comparative analysis and structure-prediction workflows.

## Methods

### The Riboseek algorithm

#### Alphabet

Riboseek converts nucleotide sequences into overlapping di-mer sequences, in which each pair of consecutive nucleotides is represented as a single di-mer token using a stride of one nucleotide. Because no complete di-mer can be formed at the final position, and because input sequences may contain ambiguous or noncanonical nucleotides, Riboseek uses eight wildcard di-mer tokens in addition to the 16 canonical di-mers. The mapping operates identically on thymine-containing dimers and uracil-containing dimers, making Riboseek applicable to both RNA and DNA.

#### Di-mer substitution matrix

Using the Rfam-derived training set alignments (see “Rfam-derived datasets: Training set for matrix estimation”) we estimated the di-mer substitution matrix as log-odds scores,

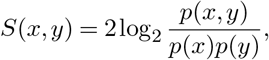

where *x* and *y* denote di-mer tokens aligned in the same alignment column, *p*(*x, y*) is the observed substitution frequency, and *p*(*x*) and *p*(*y*) are the corresponding background probabilities.

#### Riboseek E-values

To fit the location *µ* and scale *λ* parameters of the Extreme Value Distribution, we randomly sampled 100 Rfam families from the Rfam-derived training set (see section) and randomly selected one sequence from each family as a query sequence. We generated raw null hit scores by randomly shuffling each query 10,000 times and aligning it to the original query using Riboseek. We fitted 100 *µ* and *λ* parameter pairs from the null hits of each query, using the Gumbel maximum-likelihood fitting function from `hmmcalibrate` in HMMER 3.4 (29). We then computed the mean of the fitted parameters across the 100 queries to obtain: *µ* = 36.332 and *λ* = 0.185. Next, we performed post hoc calibration by first sampling 10,000 additional sequences from the training set using `seqkit sample` with `-n 10,000` (30) and generating 5,000 randomly shuffled sequences for each sampled sequence. Each shuffled sequence was aligned to its original sequence (null hit), and its E-value was computed based on the estimated *µ* and *λ*. Then, for each theoretical E-value threshold, we computed the fraction of null hits that received a smaller-than or equal-to parameter-based E-value, resulting in:

Riboseek E-value = 0.038 (parameter-based E-value)^0.828^

#### E-value benchmark

To evaluate the calibration independently, we sampled 9,857 query sequences from RNAcentral using `seqkit sample`, excluding sequences identical to those used for Riboseek E-value fitting. To produce null hits, we generated 10,000 randomly shuffled sequences from the query. We then exhaustively searched all query sequences against the shuffled sequences and compared the reported E-values to the theoretical E-value distribution expected under the null.

#### Profile construction

Before searching, Riboseek uses the di-mer substitution matrix to compute the query’s position-specific scoring matrix (PSSM), which serves as its profile. To account for the reverse-complement strand, it generates a second profile in which, for each query-side di-mer, the substitution score assigned to each target-side di-mer is replaced by the score assigned to its reverse complement. For example, the score for the AA-to-AC substitution is replaced by that for the AA-to-GU substitution.

#### Alphabet ablation

To attribute sensitivity gains to individual components, we compared five scoring schemes: a mononucleotide alphabet, the Riboseek di-mer substitution matrix without composition-bias correction, the same matrix with composition-bias correction, and the di-mer matrix of BlastR at both its default gap penalties and its best-performing penalties. All schemes were evaluated with exhaustive Smith– Waterman alignment, without prefiltering, low-complexity masking or iterative search, so that differences reflect the scoring model alone. Except for the BlastR default configuration, gap-opening and gap-extension penalties were optimised independently for each scheme by grid search over the ranges used for Riboseek (see “Parameter optimization”), and the best-performing configuration is reported; the optimum for the BlastR matrix was a gap-opening penalty of 11 and a gap-extension penalty of 1. Sensitivity was measured as mean ROC1-AUC over the validation-set queries following “Rfam benchmark”. Mean ROC1-AUC values were 0.286 (mononucleotide), 0.267 (BlastR, default penalties), 0.298 (BlastR, optimised penalties), 0.298 (Riboseek di-mer without composition-bias correction) and 0.331 (Riboseek di-mer with composition-bias correction) (Supplementary Fig. 1).

#### Search

By utilizing MMseqs2 modules, Riboseek first reduces the search space by prefiltering the target database with either the ungapped prefilter (with optional GPU acceleration) or the *k*-mer-based prefilter. It then applies the Smith– Waterman (SW) algorithm to the prefiltered targets to obtain their optimal local pairwise alignment with the query.

#### Parameter optimization

Riboseek uses two hyperparameters for pseudocounts when computing the PSSM: pca and pcb, one hyperparameter for composition bias correction: the window size, and two SW hyperparameters: the gap-opening and gap-extension penalties. The values of these parameters were selected by a grid search over pca values from 1.0 to 1.5 (step size = 0.1), pcb values from 1.3 to 2.0 (step size = 0.1), window size from 30 to 100 (step size = 5), gap-opening penalties from 13 to 25 (step size = 1), and gap-extension penalties from 1 to 3 (step size = 1). Each parameter combination was evaluated on a validation set (see “Rfam-derived datasets: Validation and test sets”), choosing the configuration under which the 61 validation queries had the highest average ROC1-AUC score (search and scoring logic described in: “Rfam benchmark”). The selected values for pca, pcb, the window size, the gap-opening penalty, and the gap-extension penalty were 1.1, 1.8, 75, 23, and 1, respectively.

#### Multiple sequence alignment

For iterative searches, Riboseek constructs a di-mer profile from the query-centered multiple sequence alignment (MSA) generated from the SW alignments and uses it in subsequent search rounds.

#### Covariance model refinement

Optionally, users may construct a covariance model (CM) from the Riboseek alignments together with a secondary structure predicted by RNAfold from the ViennaRNA package. The resulting CM can be used to realign hits with the CYK algorithm. To reduce runtime, re-alignment may be restricted to a user-defined window around the initially aligned region.

### Rfam-derived datasets

#### Initial partitioning

We obtained the 4,170 seed alignments for all families in Rfam 14.10 and constructed an initial training, validation, and test partition as follows. We first reduced redundancy by filtering out sequences in each seed alignment using the command `mmseqs filtera3m` from MMseqs2 commit `17.b804f` with the `--max-seq-id 0.8` option. After this step, 132 families had more than 20 members in their seed alignment (“big” families) and the rest (“small”) had fewer. We randomly assigned all “big” families to the validation and test partitions. To obtain an approximately 80:10:10 split, we augmented the validation and test partitions with randomly selected “small” families, assigning the remaining families to the training partition. We then enforced clan-level separation between the data partitions: if a clan in the test partition had members in another partition, those families were moved to the test partition; similarly, if the validation and training partitions shared a clan, the corresponding families were moved to the validation partition. This initial split resulted in 3,235 (78% of all families, all “small”), 472 (11%; 61 “big” + 411 “small”), and 463 (11%; 71 “big” + 392 “small”) families in the training, validation, and test partitions, respectively. The datasets used in the experiments were obtained by further processing these initial partitions.

#### Training set for matrix estimation

Starting with the initial training partition, we retrieved the original seed alignments (before redundancy reduction) and collected for each its un-aligned sequences. The unaligned sequences of each family were then clustered using `mmseqs cluster` with `- c 0.75`. We then reconstructed the training alignments by retaining only those rows in the original seed alignment whose sequences had been selected as cluster representatives.

#### Validation and test sets

To reduce benchmark runtime, our final validation and test sets included 143 and 155 families, respectively, by taking all “big” families from the initial partitions and randomly selecting 82 and 84 additional “small” families out of the 411 and 392 available ones. This meant 329 and 308 “small” families from the initial partitioning were not included in any set.

### Rfam benchmark

The following describes the analysis of the Rfam-derived test set, but the same steps were applied to the validation set for parameter optimization (see section).

#### Queries

The longest sequence in the 71 Rfam-derived test set families with more than 20 members was used as query.

#### Target database with homologs and non-homologs

We constructed a single target database of 96,701 sequences, in which each sequence from the 155-family Rfam-derived test set has known homologs (all other sequences from the same family) and known non-homologs (sequences from different Rfam clans and 10 artificial sequences produced by randomly shuffling each sequence). To prevent global alignment dominance and simulate conditions of genomic search, we used the Infernal (31) profile HMMs to generate random pseudogenomic segments ranging from 100 to 1,000 nt in length, and appended them at the 5^*′*^ and 3^*′*^ ends of each sequence in the target database.

#### Measuring sensitivity

Each query was searched against the target database. True and false positives were determined according to the known homologs and non-homologs, while hits from the same Rfam clan were ignored. When multiple hits were reported for the same query–target pair, only the top-ranked hit was considered. ROC1-AUC, defined as the fraction of detected true-positive hits ranked before the first false-positive hit, was computed for each query. Precision and recall were computed from the same true- and false-positive labels across score thresholds.

#### Tool commands

##### nhmmer

`HMMER` 3.4 was run as:

~~~
nhmmer -A nhmmer_out.sto --tblout nhmmer_out.m8
-E 10000000 query.fasta target.fasta
~~~

For iterative search, we built an HMM from the MSA reported by `nhmmer` using `hmmbuild`, and then searched the target database again using the resulting HMM as the query instead of `query.fasta`:

~~~
hmmbuild nhmmer_it.hmm nhmmer_out.sto
nhmmer -A nhmmer_iter.sto --tblout nhmmer_iter.m8
-E 10000000 nhmmer_it.hmm target.fasta
~~~

##### Infernal

`Infernal` 1.1.5 was run as:

~~~
cmbuild query.cm query.sto
cmsearch -A infernal_out.sto --tblout infernal.m8
-E 10000000 query.cm target.fasta
~~~

Here, `RNAfold` v2.7.0 was used to predict the secondary structure of each query, and each query sequence together with its predicted secondary structure was formatted as a Stock-holm file, `query.sto`. For iterative search, we built a CM from the MSA reported by `Infernal` using `cmbuild`, and then searched the target database again using the resulting CM as the query instead of `query.cm`:

~~~
cmbuild infernal_it.cm infernal_out.sto
cmsearch -A infernal_iter.sto --tblout infernal.m8
-E 10000000 infernal_it.cm target.fasta
~~~

##### BLASTN

`blastn` 2.16.0+ was run as:

~~~
blastn -task blastn -db target -query query.fasta
-out blastn.m8 -evalue 10000 -outfmt 6
~~~

##### Smith–Waterman (SW)

To assess a brute-force local-alignment baseline, we ran SSW (32) with a gap-opening penalty of 7, a gap-extension penalty of 1, a minimum reported alignment score of 1, and reverse-complement alignment enabled. The command was run as follows:

~~~
ssw_text -o 7 -e 1 -f 1 -r target.fasta query.fasta
~~~

##### Riboseek

Riboseek commit bc2a052 was run with the ungapped prefilter as follows:

~~~
riboseek search queryDB targetDB resultDB tmp
--prefilter-mode 1 -e 100000 --gpu 1
~~~

For the CPU-only configuration, the *k*-mer prefilter was used by removing the `--gpu` and `--prefilter-mode` 1 flag.

### Dfam benchmark

#### Dfam-derived test set

Starting from 4,149 Dfam 2.0 families, we first selected 1,582 families with literature support: a non-empty “Reference Location” entry. We then ran `mmseqs filtera3m` with `--max-seq-id 0.8` and retained 626 families with more than 20 members after redundancy reduction. To reduce computational cost, we randomly sampled 300 sequences from families that had more members, while ensuring that the longest sequence was included and served as the family’s representative. We then clustered all family representatives using `mmseqs cluster` with `-c 0.75`, keeping 560 families whose representative was also a cluster representative. From them, 150 families were sampled for the test set.

#### Target databases with homologs and non-homologs

We constructed a separate target database for each of the 150 families, where family members serve as known homologs of each other, and ten randomly shuffled sequences were produced for each family member, serving as artificial non-homologs. Because mononucleotide shuffling destroys dinucleotide composition and could favour a di-mer-based method, we repeated the benchmark with decoys generated by dinucleotide-preserving shuffling. Mean ROC1-AUC was 0.323 for Riboseek, 0.292 for nhmmer, and 0.237 for BLASTN, preserving the ranking obtained with mononucleotide-shuffled decoys.

#### Querying and measuring sensitivity

The longest sequence in each of the 150 Dfam-derived test set was queried against its family target database. True and false positives were determined according to the known homologs and non-homologs. When multiple hits were reported for the same query–target pair, only the top-ranked hit was considered. ROC1-AUC was computed for each query as for Rfam (see section).

#### Tool commands

`Riboseek, nhmmer SW`, and `BLASTN` were run using the same commands as in the Rfam benchmark (see section). Here, the parameters of `SW` were modified to a gap-opening penalty of 6 and a gap-extension penalty of 2, and `BLASTN` was run with `-dust no -soft_masking false`, which disables masking because Dfam families often contain low-complexity regions.

### Secondary structure prediction benchmark

#### Test set

We obtained 361 non-redundant RNA chains used for secondary-structure prediction in rMSA and extracted reference base pairs using DSSR (33). We ran rMSA for up to 30 days and performed the benchmark using the 346 RNA MSAs (Supplementary Table 3) that had been completed within this time limit.

#### Search and structure detection configurations

All tools searched against the RNAcentral v26 and NT (downloaded on 13th January 2026) databases. Each resulting MSA was de-duplicated by keeping only a single copy of duplicate sequences. We used two methods to detect conserved RNA structures in each MSA: R-scape 2.0.4.a (34) was run with default parameters, and plmc commit `18c9e55` (35) was run with `-a .ACGU -le 20.0 -lh 0.01 -m 1000`. When multiple predicted base pairs shared the same nucleotide position, only the top-ranked pair was retained. Only Watson–Crick and wobble base pairs with a sequence separation greater than three nucleotides were considered in the benchmark.

#### Measuring accuracy

Predicted base pairs were compared against the reference structure pairs for each sequence. F1 scores were computed as the harmonic mean of precision and recall: *F*_1_ = 2(precision *×* recall)*/*(precision + recall), where precision is the fraction of predicted pairings found in the reference (TP / (TP + FP)) and recall is the fraction of reference pairings that were correctly recovered (TP / P).

#### Tool-specific configurations

##### rMSA

We ran rMSA commit 0d2f660 with default parameters, with one modification. rMSA normally first runs `cmscan` to constrain the search space using covariance models available in Rfam. Because most of our test queries belong to families already curated in Rfam, this step would give rMSA access to a manually built family model that no other tool receives, so we disabled it.

##### nhmmer

We used the same pipeline as AlphaFold to construct MSAs with nhmmer. Specifically, each query sequence was searched against each database using `nhmmer` with `-E 0.001`, except that `–F3 0.02` was used for queries shorter than 50 nt. Up to 30,000 sequences were retained from each database search. The hits were realigned using `hmmalign` with a profile HMM constructed by `hmmbuild`. The two MSAs generated from the two databases were then merged after removing identical sequences.

##### Riboseek

Riboseek searched RNAcentral for three iterations (`--num-iterations 3`) with an inclusion and reporting threshold of -e 0.1 and at most 30,000 retained targets per query (`--max-seqs 30000`) per database or search iteration. A profile was built from the resulting hits with `riboseek result2profile` (`--e-profile 0.1`) and used for a single search iteration against NT with the same settings.

Both searches used the GPU-accelerated ungapped prefilter (`--prefilter-mode 1 --gpu 1`). MSAs were generated from the combined RNAcentral and NT hits and passed to R-scape or plmc.

For the structure-aware variant (Riboseek + cmsearch), a covariance-model database was built from the combined hits of both searches with `riboseek cmbuild` (`--cmlite-msa-eval 1e-3`), and hits were realigned with `riboseek cmsearch` using a window of three times the initially aligned region (`--cm-region 3.0`) and no E-value cut-off (`-e inf`).

### Tertiary structure prediction benchmark

We constructed two tertiary structure prediction benchmarks, as follows.

#### PDB-derived benchmark

All RCSB PDB (36) entries containing at least one RNA polymer entity were retrieved using the RCSB Search API and split by initial release date. Entries released before 2022 served as the reference set, whereas entries released from 2022 onward formed the query set. This temporal split resembles those used to benchmark AlphaFold 3 and Protenix-v2.

RNA chains of both the reference and the query sets were extracted from mmCIF files using GEMMI 0.7.4 (37). For multi-model entries, only the first model was retained. Non-RNA chains and short RNA fragments containing fewer than 10 nucleotides were removed. To retain RNA chains with limited protein interactions and well-defined intramolecular structure, we kept only chains for which fewer than 30% of the residues contacted protein atoms, defined as heavy atoms within 4 Å, and more than 90% of the residues formed intramolecular contacts. We also excluded RNA chains from large complexes containing 10 or more chains.

Each query RNA chain was compared to all reference RNA chains using US-align (38) in nucleic-acid mode, and the TM-score normalized by the shorter chain length was recorded as a symmetric measure of shared substructure. Query chains were classified as structurally similar to known RNAs if their best TM-score against any reference chain exceeded 0.45, which has been reported as a threshold for RNAs from the same family (39), and as structurally novel otherwise. This resulted in 411 similar chains and 109 novel chains. To remove sequence redundancy, we excluded chains with greater than 80% sequence identity to any reference chain, as reported by US-align. The remaining query chains were then aligned to each other in an all-against-all manner.

We constructed a graph by connecting pairs of chains with TM-scores greater than 0.45 and clustered the graph using the Markov Cluster Algorithm (MCL) (40) with an inflation parameter of 2.0. For each cluster, the chain with the highest mean TM-score to all other cluster members was selected as the representative. Finally, to focus the benchmark on less common natural RNA classes, ribosomal RNA, transfer RNA, and synthetic RNA chains were excluded. This procedure resulted in 13 similar (training-like) RNA chains and 15 novel RNA chains (Supplementary Table 4, 5).

MSAs for training-like and novel RNA chains were generated using `nhmmer`, `rMSA`, and `Riboseek`, and were then provided as input to `AlphaFold3` and `Protenix-v2` for tertiary structure prediction. The predicted structures were evaluated by computing the TM-score, normalized by the length of the reference PDB chain, using `US-align`, and the lDDT using `OpenStructure` (41).

#### Tool configurations

##### AlphaFold 3

AlphaFold 3 v3.0.0 was run with random seeds 10 and 42, and the model with the highest ranking score was retained.

##### Protenix-v2

Protenix-v2 was run with random seed 101 with custom RNA MSA input enabled using `--use_rna_msa True --use_msa True`, and `protenix-v2` was used as the model weights.

##### US-align

US-align version 20241108 was run with `-mol RNA -TMscore 1`, and the TM-score normalized by the length of the reference PDB structure was reported.

##### OpenStructure

OpenStructure 2.11.0 was run with `–rna --lddt --local-lddt --lddt-no-stereochecks`, and the mean lDDT score was reported.

### Precomputed RNA MSA resource

We downloaded all sequences from RNAcentral Release 25 and removed sequences containing gap characters or nonstandard nucleotides, followed by exact-sequence deduplication. Secondary structures were predicted with EternaFold(42) using default settings. We retained sequences for which at least 60% of nucleotides were predicted to participate in base pairing and excluded sequences of 600 nucleotides or longer, resulting in 1,731,677 query sequences. Riboseek was then used to generate one MSA for each retained sequence, following the same two-stage procedure as in the secondary-structure benchmark (see “Tool-specific configurations”) without searching the NT database.

For comparison, we obtained the RNAcentral and Rfam MSA sets released with OpenFold3. MSA depth was defined as the number of sequences in an alignment. Effective sequence counts (*N*_eff_) were computed with hhmake (43), and length-normalized diversity was calculated as 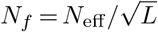, where *L* is the query length. The paired comparison with the Open-Fold3 RNAcentral set was restricted to 4,991 queries present in both collections.

### Scalability benchmark

For the search-time benchmark, we used 155 sequences from the Rfam-derived test set and replaced the target database with species-specific sequence sets from RNAcentral v26.

For the MSA-generation time benchmark, we randomly sampled seven queries with varying sequence lengths. All tools searched against RNAcentral v26 and NT. To measure batch throughput at a realistic compute-to-I/O ratio, the seven queries were replicated tenfold into a single 70-query database and searched in one Riboseek run, since Riboseek parallelises across queries. The other tools parallelise across target sequences and rescan the full database for each query; they were run on each of the seven queries individually, and their combined runtimes were obtained by multiplying the summed per-query runtimes by ten. Because NT is too large to remain in 512 GB of RAM between queries, runtime is linear in query count, so this extrapolation approximates a direct 70-query run.

### Computing resources and runtime measurement

All runtimes reported in this work were measured on the same two servers, each with 512 GB of RAM and target databases stored on local NVMe disks. Riboseek was run on a single-socket Intel Xeon Platinum 8559C (24 cores, 48 threads) with one NVIDIA RTX PRO 6000 Blackwell Server Edition GPU; nhmmer, Infernal, BLASTN, SW, rMSA and cmsearch of riboseek were run on a dual-socket Intel Xeon Gold 6530 (64 cores, 128 threads).

### Web server Web API

Riboseek is integrated into the Foldseek webserver (44) for accessible, interactive use requiring only a web browser. The server offers a database derived from RNAcentral, Rfam, and the NT transcript sequences used in AlphaFold3, clustered with MMseqs2. Additionally, we deployed Riboseek as a web API for rapid, on-demand RNA MSA generation. It returns query-centered MSAs searched against the same database and is integrated into ColabFold for AlphaFold3 predictions, avoiding the need to download and maintain the underlying databases locally.

## Data availability

The benchmark data are available via Zenodo at https://doi.org/10.5281/zenodo.21675713. The precomputed RNA MSA resource generated in this study is openly available in the Steinegger Lab AWS Open Data bucket at s3://steineggerlab/riboseek/rna_central_msa/, and over HTTPS at https://opendata.mmseqs.org/riboseek/. The resource comprises 1,731,677 alignments distributed as 1,723 zstd-compressed archives (1.96 TiB). The dataset is available under a Creative Commons Attribution license (CC-BY 4.0).

## Code availability

Riboseek is free, open-source software. The source code, ready-to-use binaries, and precomputed databases are available at github.com/steineggerlab/riboseek. The scripts used for benchmarking and plotting are available at github.com/steineggerlab/riboseek-analysis.

## Acknowledgments

We thank E. Levy Karin (ELKMO) for insightful discussions and help revising the manuscript; J. Söding and J. Belousov for helpful discussions on nucleotide alphabets; A. Vorontsov for bug fixes to the GPU implementation; C. Dallago for discussions of *plmc*; H. Lee for discussions of RNA structure prediction; and R. Bushuiev for discussions of the benchmark design. Martin Steinegger acknowledges support by the National Research Foundation of Korea (NRF) grants funded by the Korea government (MSIT) (RS-2024-00396026, RS-2025-00438101, RS-2026-25549295), the Novo Nordisk Foundation (NNF24SA0092560). Milot Mirdita acknowledges support by the National Research Foundation of Korea (NRF) grants funded by the Korea government (MSIT) (RS-2026-25515455).

## Author contributions

**Conceptualization:** S.P., M.S. **Data curation:** S.P., A.F., A.B., S.K., M.M. **Formal analysis:** S.P., K.D. **Funding acquisition:** M.S. **Investigation:** S.P., A.B. **Methodology:** S.P., M.M., M.S. **Project administration:** S.P., M.S. **Resources:** A.F., M.S. **Software:** S.P., M.M., M.S. **Supervision:** M.M., M.S. **Validation:** S.P., S.K. **Visualization:** S.P., M.S. **Writing – original draft:** S.P., K.D., M.M., M.S. **Writing – review & editing:** S.P., K.D., A.B., M.M., M.S.

## References

1. Sayers, E. W. et al. GenBank 2025 update. Nucleic Acids Research 53, D56–D61 (2025).

2. Yuan, D. et al. The European Nucleotide Archive in 2023. Nucleic Acids Research 52, D92–D97 (2024).

3. Altschul, S. F., Gish, W., Miller, W., Myers, E. W. & Lipman, D. J. Basic local alignment search tool. Journal of Molecular Biology 215, 403–410 (1990).

4. Camacho, C. et al. BLAST+: architecture and applications. BMC Bioinformatics 10, 421 (2009).

5. McGinnis, S. & Madden, T. L. BLAST: at the core of a powerful and diverse set of sequence analysis tools. Nucleic Acids Research 32, W20–W25 (2004).

6. Wheeler, T. J. & Eddy, S. R. nhmmer: DNA homology search with profile HMMs. Bioinformatics 29, 2487–2489 (2013).

7. Eddy, S. R. & Durbin, R. RNA sequence analysis using covariance models. Nucleic Acids Research 22, 2079–2088 (1994).

8. Rivas, E. & Eddy, S. R. Noncoding RNA gene detection using comparative sequence analysis. BMC Bioinformatics 2, 8 (2001).

9. Nawrocki, E. P. & Eddy, S. R. Infernal 1.1: 100-fold faster RNA homology searches. Bioinformatics 29, 2933–2935 (2013).

10. Ontiveros-Palacios, N. et al. Rfam 15: RNA families database in 2025. Nucleic Acids Research 53, D258–D267 (2025).

11. Abramson, J. et al. Accurate structure prediction of biomolecular interactions with AlphaFold 3. Nature 630, 493–500 (2024).

12. Zhang, Y. et al. Protenix-v2: Broadening the Reach of Structure Prediction and Biomolecular Design. bioRxiv 10.64898/2026.04.10.717613 (2026).

13. The RNAcentral Consortium. RNAcentral in 2026: genes and literature integration. Nucleic Acids Research 54, D303–D313 (2026).

14. Sayers, E. W. et al. Database resources of the National Center for Biotechnology Information in 2025. Nucleic Acids Research 53, D20–D29 (2025).

15. Zhang, C., Zhang, Y. & Pyle, A. M. rMSA: a sequence search and alignment algorithm to improve RNA structure modeling. Journal of Molecular Biology 435, 167904 (2023).

16. Steinegger, M. & Söding, J. MMseqs2 enables sensitive protein sequence searching for the analysis of massive data sets. Nature Biotechnology 35, 1026–1028 (2017).

17. Jumper, J. et al. Highly accurate protein structure prediction with AlphaFold. Nature 596, 583–589 (2021).

18. Mirdita, M. et al. ColabFold: making protein folding accessible to all. Nature Methods 19, 679–682 (2022).

19. Kallenborn, F. et al. GPU-accelerated homology search with MMseqs2. Nature Methods 22, 2024–2027 (2025).

20. Smith, T. F. & Waterman, M. S. Identification of common molecular subsequences. Journal of Molecular Biology 147, 195–197 (1981).

21. Gotoh, O. An improved algorithm for matching biological sequences. Journal of Molecular Biology 162, 705–708 (1982).

22. Lorenz, R. et al. ViennaRNA Package 2.0. Algorithms for Molecular Biology 6, 26 (2011).

23. Bussotti, G. et al. BlastR—fast and accurate database searches for non-coding RNAs. Nucleic Acids Research 39, 6886–6895 (2011).

24. Storer, J., Hubley, R., Rosen, J., Wheeler, T. J. & Smit, A. F. A. The Dfam community resource of transposable element families, sequence models, and genome annotations. Mobile DNA 12, 2 (2021).

25. Mariani, V., Biasini, M., Barbato, A. & Schwede, T. lDDT: a local superposition-free score for comparing protein structures and models using distance difference tests. Bioinformatics 29, 2722–2728 (2013).

26. Studer, G. et al. A fully automated benchmarking suite to compare macromolecular complexes. Nature methods 23, 387–394 (2026).

27. Zhang, Y. TM-align: a protein structure alignment algorithm based on the TM-score. Nucleic Acids Research 33, 2302–2309 (2005).

28. The OpenFold3 Team. OpenFold3-preview (2025). URL https://github.com/aqlaboratory/openfold-3.

## References

29. Eddy, S. R. Profile hidden Markov models. Bioinformatics 14, 755–763 (1998).

30. Shen, W., Le, S., Li, Y. & Hu, F. SeqKit: a cross-platform and ultrafast toolkit for FASTA/Q file manipulation. PLoS ONE 11, e0163962 (2016).

31. Nawrocki, E. P. & Eddy, S. R. Infernal 1.1: 100-fold faster RNA homology searches. Bioinformatics 29, 2933–2935 (2013).

32. Zhao, M., Lee, W.-P., Garrison, E. P. & Marth, G. T. SSW library: an SIMD Smith-Waterman C/C++ library for use in genomic applications. PLoS ONE 8, e82138 (2013).

33. Lu, X.-J., Bussemaker, H. J. & Olson, W. K. DSSR: an integrated software tool for dissecting the spatial structure of RNA. Nucleic Acids Research 43, e142–e142 (2015).

34. Rivas, E., Clements, J. & Eddy, S. R. Estimating the power of sequence covariation for detecting conserved RNA structure. Bioinformatics 36, 3072–3076 (2020).

35. Hopf, T. A. et al. Mutation effects predicted from sequence co-variation. Nature Biotechnology 35, 128–135 (2017).

36. Burley, S. K. et al. Updated resources for exploring experimentally-determined PDB structures and Computed Structure Models at the RCSB Protein Data Bank. Nucleic Acids Research 53, D564–D574 (2025).

37. Wojdyr, M. GEMMI: A library for structural biology. Journal of Open Source Software 7, 4200 (2022).

38. Zhang, C., Shine, M., Pyle, A. M. & Zhang, Y. US-align: universal structure alignments of proteins, nucleic acids, and macromolecular complexes. Nature Methods 19, 1109–1115 (2022).

39. Gong, S., Zhang, C. & Zhang, Y. RNA-align: quick and accurate alignment of RNA 3D structures based on size-independent TM-scoreRNA. Bioinformatics 35, 4459–4461 (2019).

40. Van Dongen, S. Graph clustering via a discrete uncoupling process. SIAM Journal on Matrix Analysis and Applications 30, 121–141 (2008).

41. Biasini, M. et al. OpenStructure: an integrated software framework for computational structural biology. Acta Crystallographica Section D: Biological Crystallography 69, 701–709 (2013).

42. Wayment-Steele, H. K. et al. RNA secondary structure packages evaluated and improved by high-throughput experiments. Nature Methods 19, 1234–1242 (2022).

43. Steinegger, M. et al. HH-suite3 for fast remote homology detection and deep protein annotation. BMC Bioinformatics 20, 473 (2019).

44. Mirdita, M., Steinegger, M. & Söding, J. MMseqs2 desktop and local web server app for fast, interactive sequence searches. Bioinformatics 35, 2856–2858 (2019).

