## Supplementary Materials for "Fast remote nucleotide sequence alignment with Riboseek"

July 31, 2026

### Supplementary Note 1: Supplementary Materials

#### A. Figures

- Supplementary Fig. 1. Complete Rfam benchmark result.
- Supplementary Fig. 2. Ablation study of substitution matrix and composition bias correction.
- Supplementary Fig. 3. Comparison of TM-score among different RNA structure predictors for training-like targets.
- Supplementary Fig. 4. Comparison of lDDT among different RNA structure predictors for training-like targets.
- Supplementary Fig. 5. Comparison of TM-score among different RNA structure predictors for novel targets.
- Supplementary Fig. 6. Comparison of lDDT among different RNA structure predictors for novel targets.

#### B. Tables

- Supplementary Table 1. Performance of RNA structure prediction methods using different MSA generation approaches on a training-like set.
- Supplementary Table 2. Performance of RNA structure prediction methods using different MSA generation approaches on a novel set.
- Supplementary Table 3. Detailed list of 346 RNA PDB IDs for secondary structure prediction.
- Supplementary Table 4. Detailed list of 13 RNA PDB IDs for training-like test set.
- Supplementary Table 5. Detailed list of 15 RNA PDB IDs for novel test set.

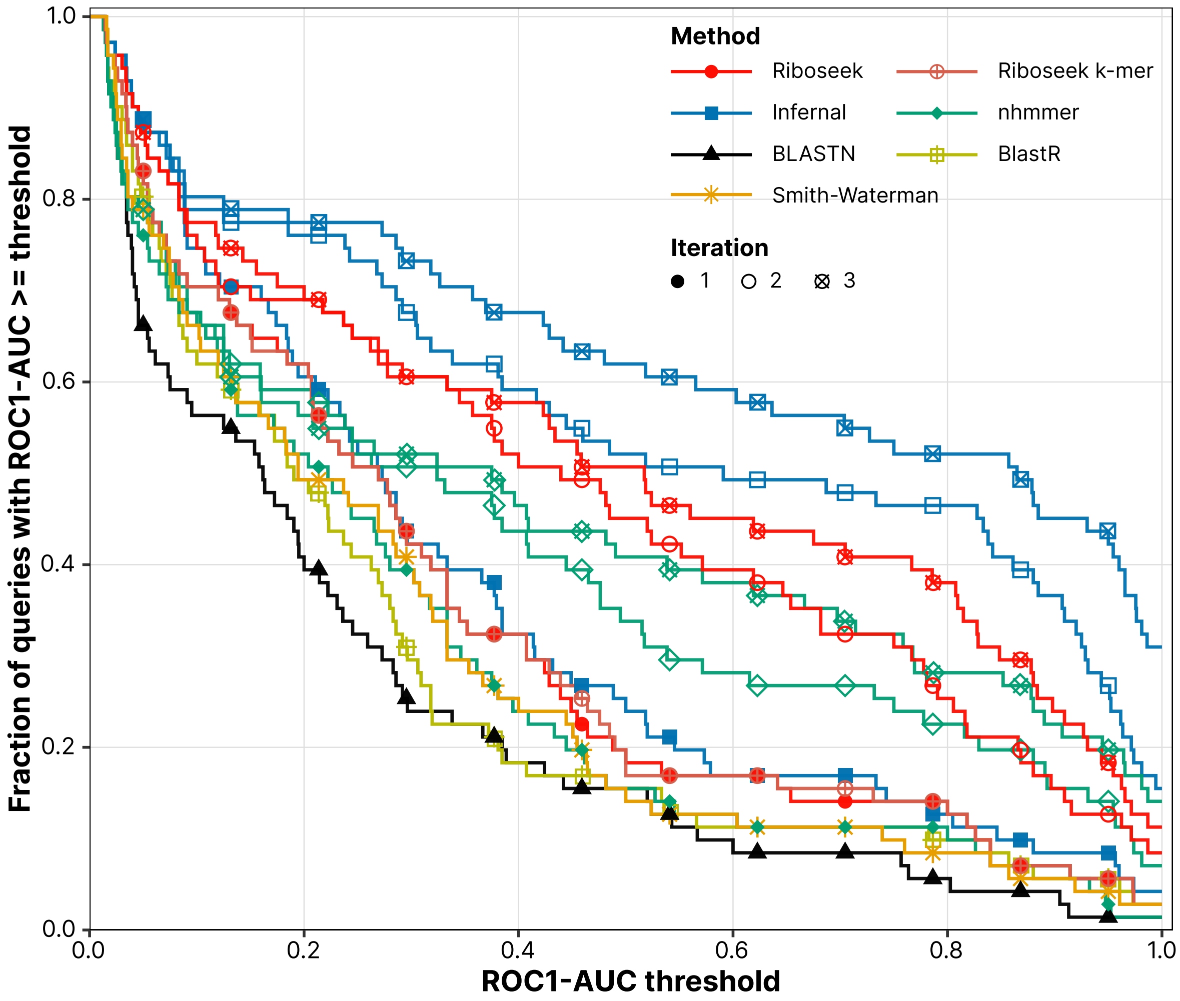


**Fig. 1.** Complete Rfam benchmark result.

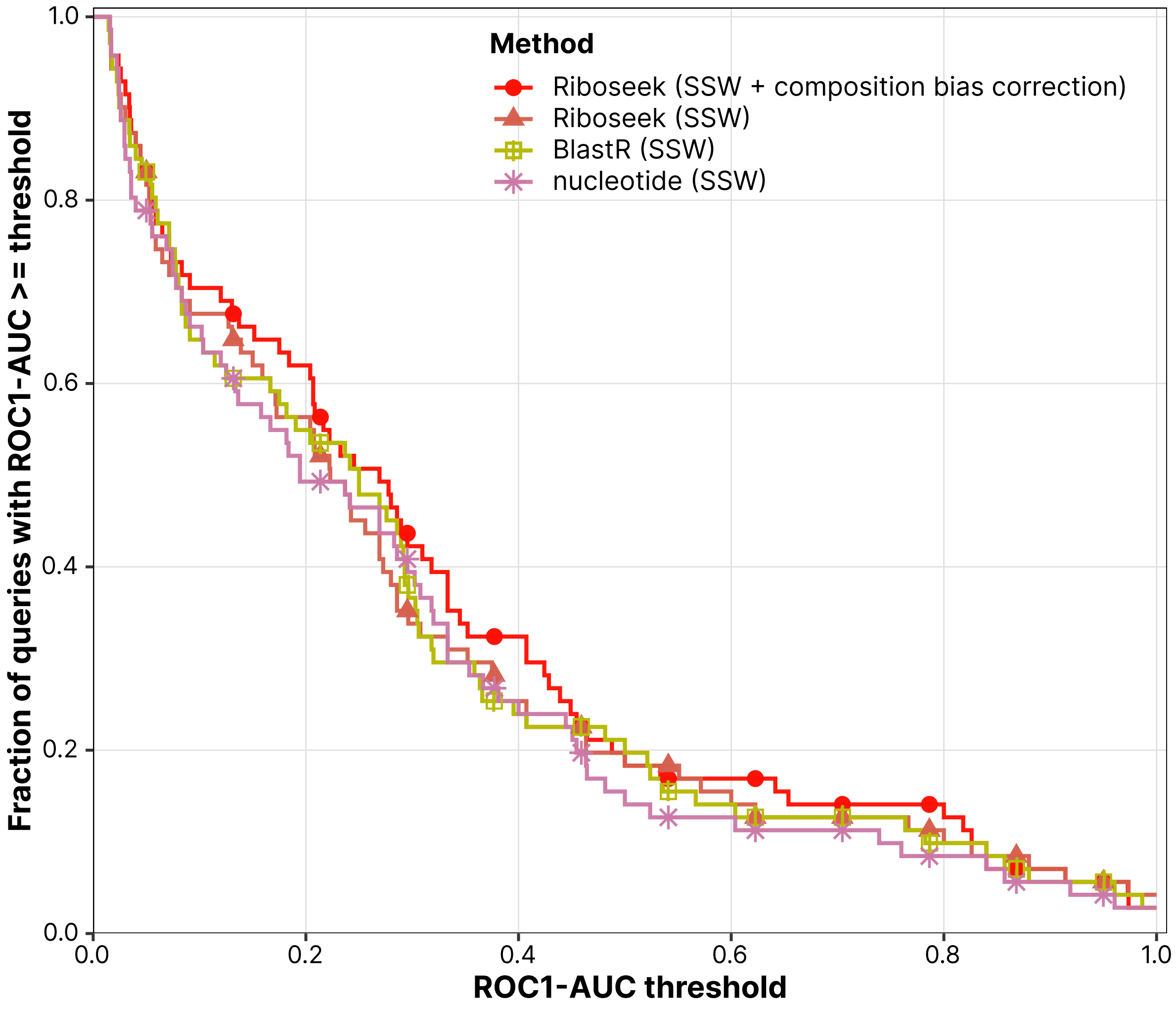


**Fig. 2.** Ablation study of substitution matrix and composition bias correction.

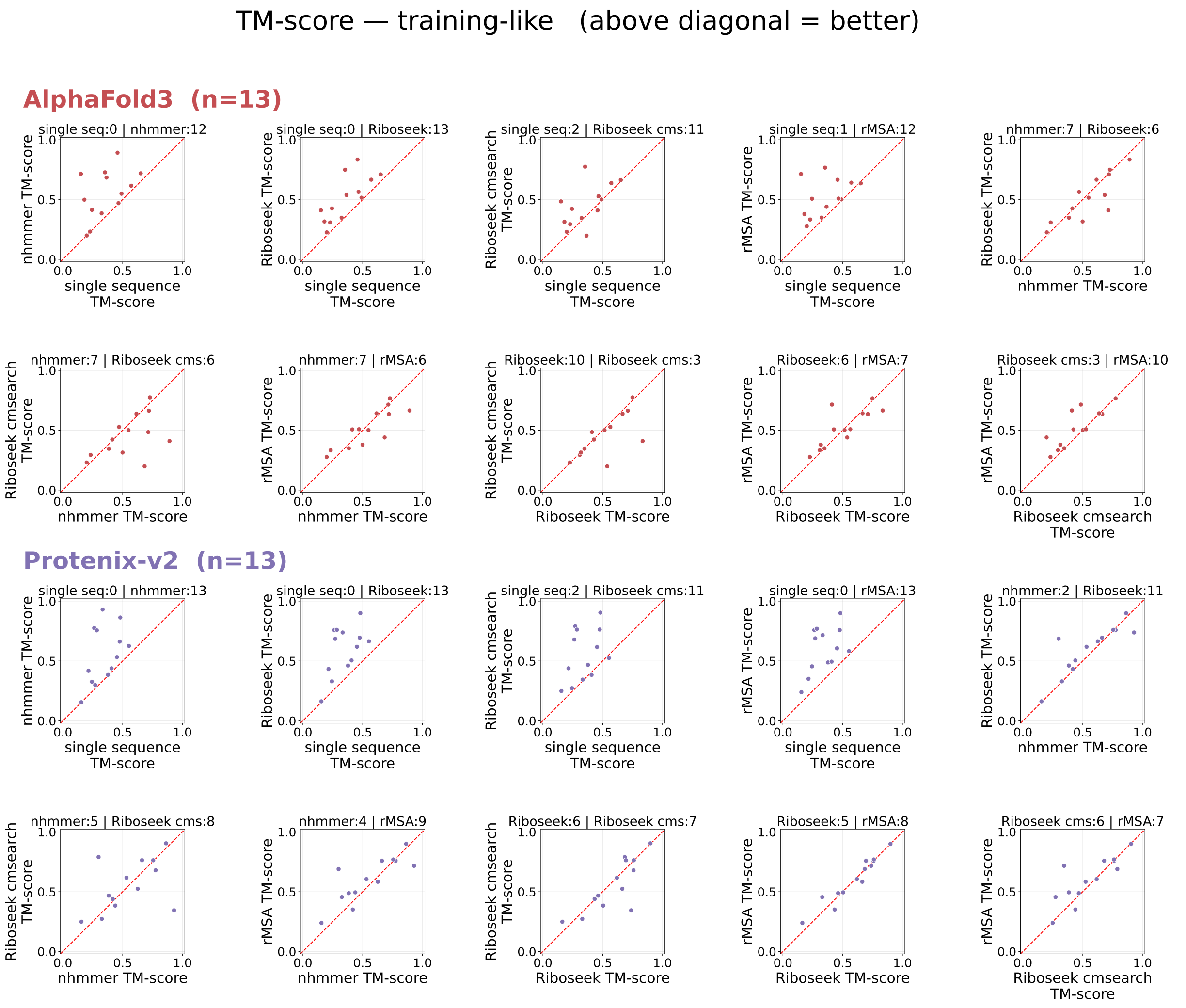


**Fig. 3.** Comparison of TM-score among different RNA structure predictors for training-like targets.

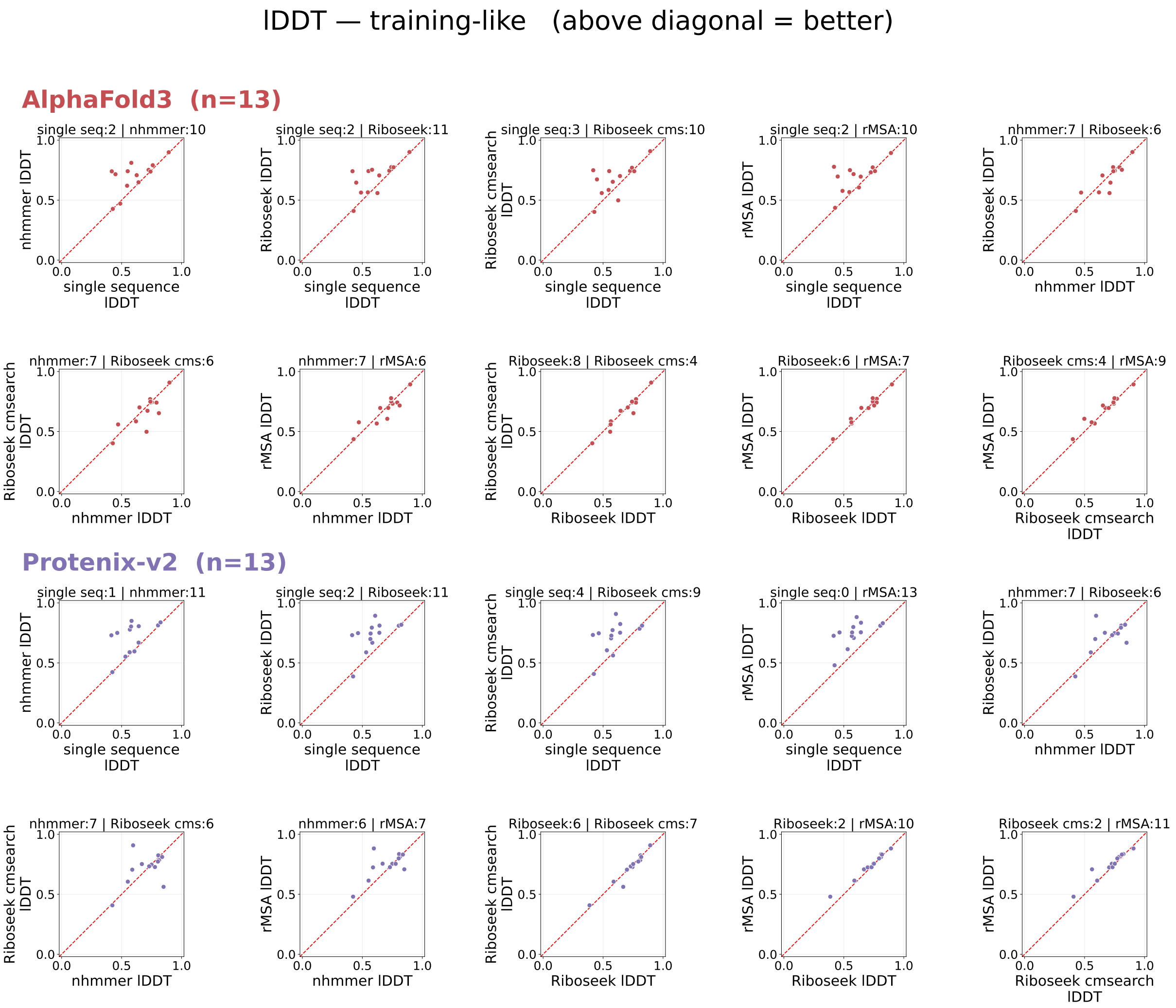


**Fig. 4.** Comparison of lDDT among different RNA structure predictors for training-like targets.

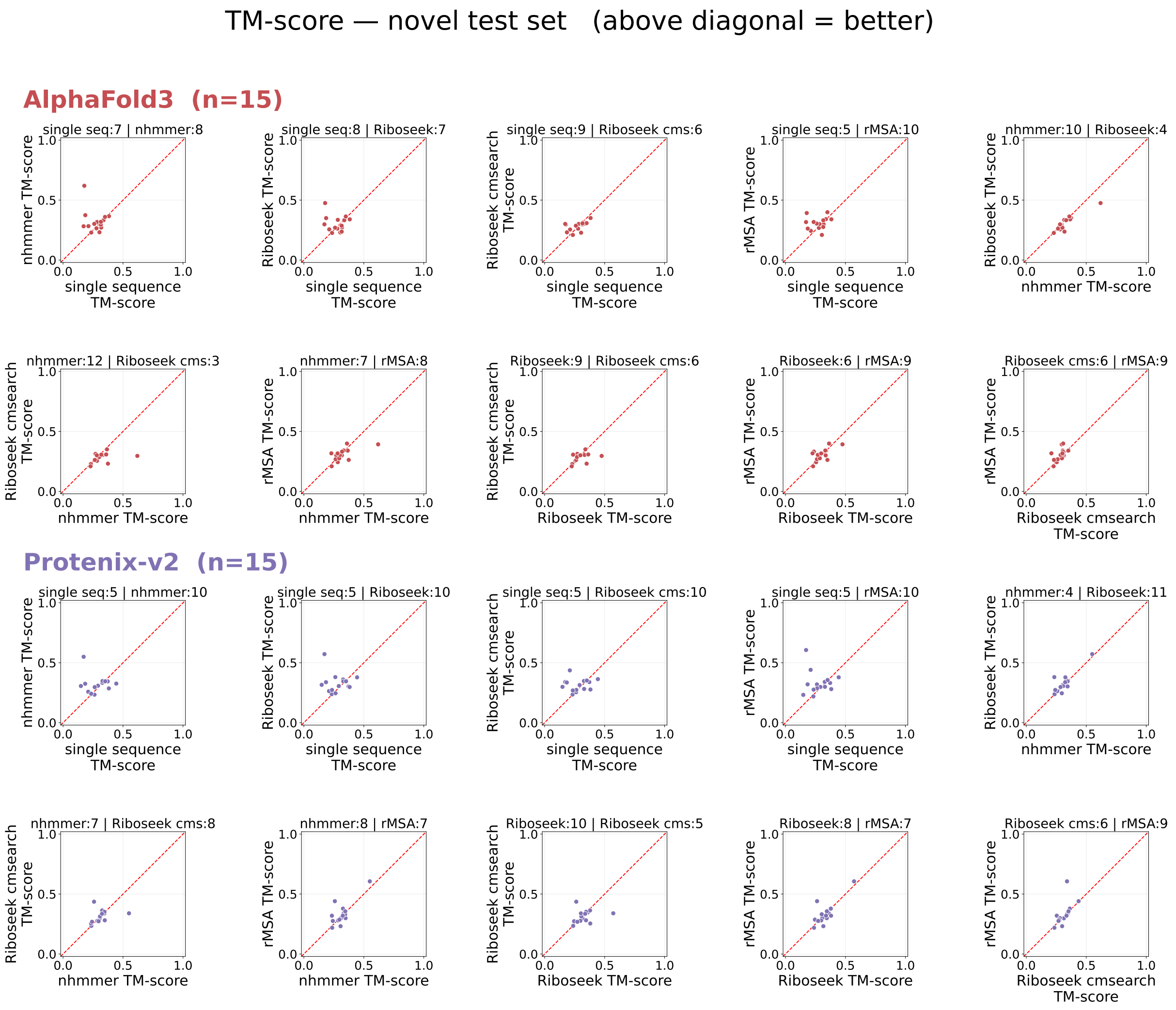


**Fig. 5.** Comparison of TM-score among different RNA structure predictors for novel targets.

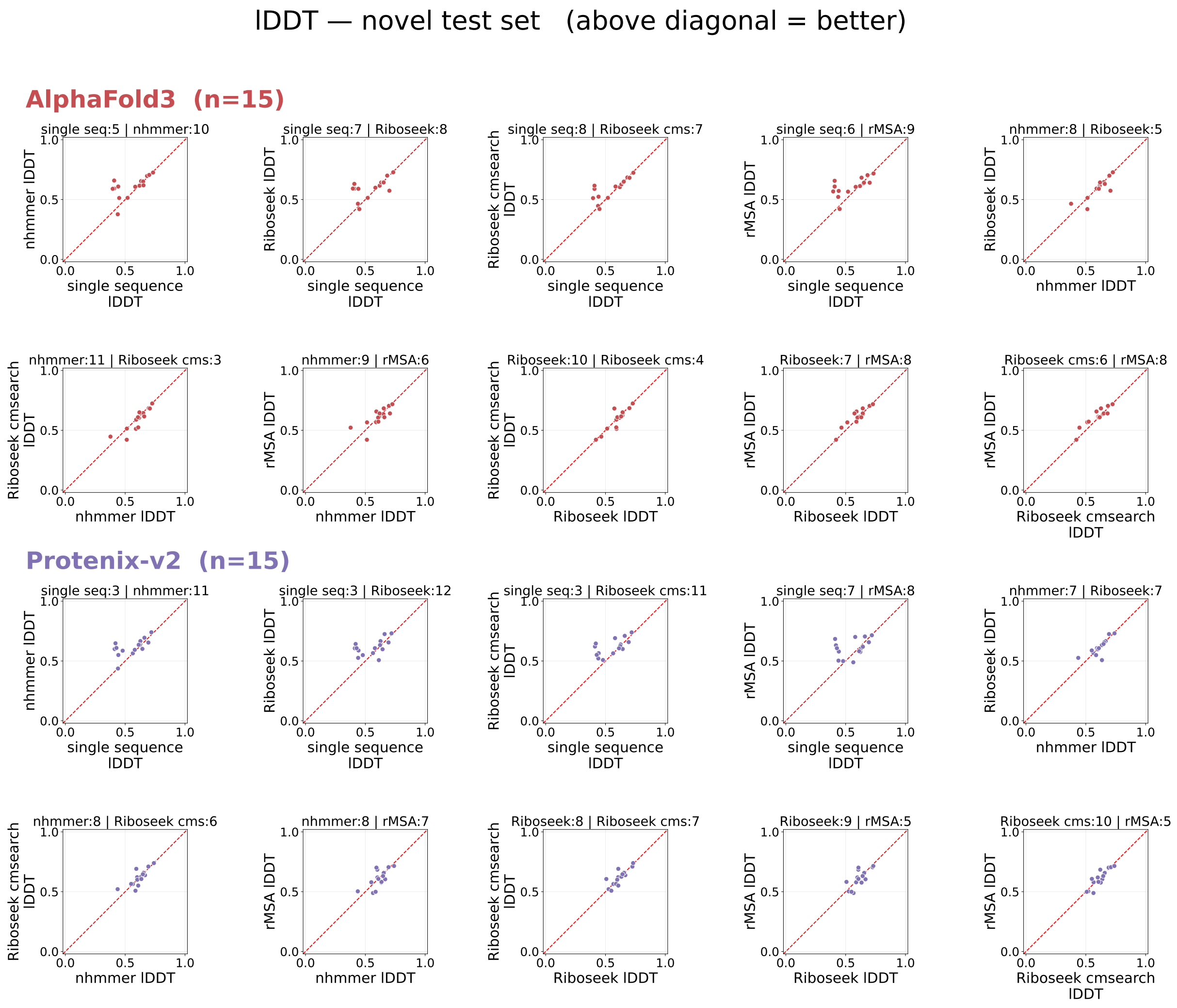


**Fig. 6.** Comparison of lDDT among different RNA structure predictors for novel targets.

Table 1. Performance of RNA structure prediction methods using different MSA generation approaches on a training-like set.

| Predictor | Metric | single sequence | nhmmer | Riboseek | Riboseek cmsearch | rMSA |
|---|---|---|---|---|---|---|
| AlphaFold3 | lDDT ↑ | 0.604 | **0.697** | 0.683 | 0.671 | 0.690 |
| | TM ↑ | 0.360 | **0.546** | 0.509 | 0.446 | 0.517 |
| Protenix-v2 | lDDT ↑ | 0.589 | 0.707 | 0.726 | 0.718 | **0.744** |
| | TM ↑ | 0.347 | 0.551 | 0.593 | 0.554 | **0.601** |

Table 2. Performance of RNA structure prediction methods using different MSA generation approaches on a novel set.

| Predictor | Metric | single sequence | nhmmer | Riboseek | Riboseek cmsearch | rMSA |
|---|---|---|---|---|---|---|
| AlphaFold3 | lDDT ↑ | 0.555 | 0.609 | 0.597 | 0.590 | **0.610** |
| | TM ↑ | 0.275 | **0.323** | 0.302 | 0.283 | 0.307 |
| Protenix-v2 | lDDT ↑ | 0.556 | 0.614 | 0.614 | **0.616** | 0.610 |
| | TM ↑ | 0.280 | 0.316 | 0.331 | 0.314 | **0.332** |

**Table 3.** Detailed list of 346 RNA PDB IDs for secondary structure prediction.

| PDB ID | Chain ID | Description |
|---|---|---|
| 1DK1 | B | rRNA fragment |
| 1ET4 | A | RNA Aptamer |
| 1F1T | A | Malachite Green Aptamer RNA |
| 1FG0 | A | 23S rRNA |
| 1G59 | B | tRNA-GLU |
| 1GAX | C | tRNA-VAL |
| 1H3E | B | tRNA-TYR(GUA) |
| 1H4S | T | tRNA-PRO (CGG) |
| 1I6U | C | 16s rRNA fragment |
| 1IL2 | C | Aspartyl Transfer RNA |
| 1J1U | B | tRNA-TYR |
| 1J2B | C | tRNA-VAL |
| 1JBR | D | SRD RNA analog |
| 1KOG | I | Threonyl-tRNA synthetase mRNA |
| 1KXK | A | ai5g group II Self-splicing intron |
| 1M5O | E | RNA Hairpin Ribozyme |
| 1MZP | B | fragment of 23S rRNA |
| 1NBS | A | Ribonuclease P RNA |
| 1P6V | B | tRNA domain of transfer-messenger RNA |
| 1QF6 | B | tRNA-THR |
| 1S03 | A | spc Operon mRNA |
| 1SER | T | tRNA-SER |
| 1U0B | A | tRNA-CYS |
| 1U63 | B | Fragment of mRNA for L1 |
| 1U6B | B | Self-Splicing Group I Intron with Both Exons |
| 1U9S | A | Ribonuclease P |
| 1UN6 | E | 5S rRNA |
| 1VQ8 | 9 | 5S rRNA |
| 1WZ2 | C | tRNA |
| 1X8W | A | Group I Intron RNA |
| 1XJR | A | s2m RNA |
| 1Y0Q | A | Group I Intron ribozyme |
| 1YLS | B | RNA Diels-Alder ribozyme |
| 1ZHO | B | mRNA |
| 2A64 | A | ribonuclease P RNA |
| 2B63 | R | RNA inhibitor of RNA polymerase II |
| 2CSX | C | tRNA-MET |
| 2CZJ | B | tmRNA |
| 2D6F | E | tRNA |
| 2DER | C | tRNA |
| 2DLC | Y | tRNA |
| 2DR2 | B | tRNA-TRP |
| 2DU3 | D | tRNA |
| 2FK6 | R | tRNA-THR |
| 2HVY | E | H/ACA RNA from RNA pseudouridine synthases |
| 2IL9 | M | Ribosomal binding domain of IRES RNA |
| 2IY5 | T | tRNA-PHE |
| 2NRE | F | tRNA-LEU |
| 2NUE | C | RNA |
| 2OIU | P | L1 Ribozyme RNA Ligase |
| 2PXD | B | 4.5 S RNA |
| 2QUS | A | Hammerhead ribozyme |
| 2QWY | A | SAM-II riboswitch |
| 2XDB | G | TOXI |
| 2XXA | F | 4.5S RNA |
| 2YGH | A | SAM-I riboswitch |
| 2Z75 | B | glmS ribozyme RNA |
| 2ZH3 | B | tRNA |
| 2ZNI | C | Bacterial tRNA |
| 2ZUE | B | tRNA-ARG |
| 2ZZM | B | tRNA-LEU |
| 3A2K | C | bacterial tRNA |
| 3ADB | C | selenocysteine tRNA |
| 3AKZ | E | tRNA-GLN |
| 3AM1 | B | ASL-truncated tRNA |
| 3AMT | B | tRNA-ILE |
| 3BWP | A | Group IIC intron |
| 3CUL | D | Aminoacyl-tRNA synthetase ribozyme |
| 3D2V | A | TPP-specific riboswitch |
| 3DIL | A | Lysine riboswitch |

**Table 3.** (Continued.)

| PDB ID | Chain ID | Description |
|---|---|---|
| 3E5C | A | SMK box (SAM-III) Riboswitch |
| 3EGZ | B | Tetracycline aptamer and artificial riboswitch |
| 3EPH | E | tRNA |
| 3F2Q | X | FMN riboswitch |
| 3FU2 | A | PreQ1 riboswitch |
| 3GCA | A | PreQ1 riboswitch |
| 3GS5 | C | RNA |
| 3HHN | C | Class I ligase ribozyme, self-ligation product |
| 3IAB | R | P3 domain of the RNA component of RNase MRP |
| 3ICQ | D | tRNA |
| 3IVN | A | A-riboswitch |
| 3IWN | A | C-di-GMP riboswitch |
| 3J79 | B | 5S rRNA |
| 3J79 | C | 5.8S rRNA |
| 3J7A | 7 | tRNA |
| 3J7Y | B | mt-tRNA-VAL |
| 3JB9 | P | U2 snRNA |
| 3JB9 | C | U5 snRNA |
| 3JCS | 3 | 26S gamma rRNA |
| 3JCS | 8 | 5S ribosomal RNA |
| 3K0J | E | ThiM riboswitch |
| 3KTW | C | SRP RNA |
| 3L3C | P | GLMS Ribozyme |
| 3LA5 | A | Adenosine Riboswitch |
| 3LWR | D | H/ACA RNA |
| 3NDB | M | SRP RNA |
| 3NKB | B | Hepatitis delta virus ribozyme |
| 3NPQ | A | S-Adenosylhomocysteine Riboswitch |
| 3OVB | C | tRNA mimic |
| 3OWI | B | Domain II of glycine riboswitch |
| 3P22 | A | Core ENE hairpin from Kaposi's sarcoma-associated herpesviru PAN RNA |
| 3P49 | A | Glycine Riboswitch |
| 3PDR | X | M-box Riboswitch RNA |
| 3Q1Q | B | RNase P RNA |
| 3Q1Q | C | tRNA-PHE |
| 3Q3Z | V | c-di-GMP-II riboswitch |
| 3R4F | A | pRNA |
| 3RG5 | A | tRNA-SEC |
| 3RW6 | F | Constitutive transport element of Mason-Pfizer monkey virus RNA |
| 3SKI | B | 2'-Deoxyguanosine riboswitch |
| 3SNP | C | Ferritin H IRE RNA |
| 3SUH | X | Riboswitch |
| 3UMY | B | 23S rRNA |
| 3V7E | C | SAM-I riboswitch aptamer with an engineered helix P3 |
| 3W3S | B | selenocysteine tRNA |
| 3WQY | C | tRNA-ALA |
| 3ZP8 | A | Hammerhead Ribozyme, Enzyme Strand |
| 4ATO | G | TOXI |
| 4C7O | E | SRP RNA |
| 4ENC | A | Fluoride riboswitch |
| 4FE5 | B | xpt-pbuX guanine riboswitch aptamer domain |
| 4FRN | A | Cobalamin riboswitch aptamer domain |
| 4GMA | Z | Adenosylcobalamin riboswitch |
| 4GXY | A | Adenosylcobalamin riboswitch |
| 4JF2 | A | PreQ1-II Riboswitch |
| 4K27 | U | Myotonic Dystrophy Type 2 RNA |
| 4KQY | A | YitJ S box/SAM-I riboswitch |
| 4KR6 | C | Truncated tRNA |
| 4LCK | B | tRNA-GLY |
| 4LCK | C | T-box riboswitch stem I |
| 4LVW | A | THF riboswitch |
| 4M4O | B | Aptamer minE |
| 4MGN | A | glyQS T box riboswitch |
| 4O26 | E | Telomerase TR |
| 4OJI | A | Twister Ribozyme |
| 4OOG | D | RNase III cleavage product |
| 4OQU | A | SAM-I/IV riboswitch |
| 4PDB | I | SELEX RNA aptamer |
| 4PKD | V | U1 snRNA stem-loops 1 and 2 |
| 4PMI | A | Rev-Response-Element RNA |

**Table 3.** (Continued.)

| PDB ID | Chain ID | Description |
|---|---|---|
| 4PQV | A | XRN1-resistant flaviviral RNA |
| 4PRF | B | Hepatitis Delta virus ribozyme |
| 4QJD | B | Twister RNA sequence |
| 4QJH | B | Twister Ribozyme |
| 4QK9 | A | C-di-AMP riboswitch |
| 4QLN | A | ydaO riboswitch |
| 4RDX | C | tRNA-HIS |
| 4RGE | A | env22 twister ribozyme |
| 4RMO | B | Antitoxin for CptIN Type III Toxin |
| 4RUM | A | NiCo riboswitch RNA |
| 4RZD | A | PreQ1-III Riboswitch (Class 3) |
| 4UYK | R | SRP RNA |
| 4V2S | Q | Bacterial small RNAs (sRNAs) rydC |
| 4V5G | AY | A-site tRNA-THR |
| 4V5L | AY | A-site tRNA G24A tRNA-TRP |
| 4V7L | AY | tRNA-GLN |
| 4V83 | AV | domain 3 of PSIC IGR IRES RNA |
| 4V8B | AB | tRNA-LEU |
| 4V8D | AB | tRNA-TYR |
| 4V8N | AW | A-site tRNA ILE2 Agmatidine |
| 4V8P | B2 | 5.8S rRNA |
| 4V8P | B3 | 5S rRNA |
| 4V9I | AY | A-site tRNA |
| 4W90 | C | Riboswitch a pseudo-dimeric RNA |
| 4WFL | A | Bacterial SRP Alu domain |
| 4WJ4 | B | 76mer-tRNA |
| 4WZJ | V | U4 small nuclear RNA variant |
| 4X4P | B | G70A tRNA minihelix ending in CCAC |
| 4XWF | A | pfl RNA |
| 4Y1J | A | yybP-ykoY riboswitch |
| 4Y1M | A | yybP-ykoY riboswitch |
| 4YAZ | R | 3’,3’-cGAMP riboswitch |
| 4YBB | CB | 5S rRNA |
| 4YYE | C | tRNA |
| 4ZNP | A | pfI riboswitch |
| 5AH5 | C | tRNA-LEU TAA isoacceptor |
| 5AOX | C | Alu Jo consensus RNA |
| 5B2O | B | Guide RNA |
| 5B63 | B | tRNA-ARG |
| 5BJO | E | RNA aptamer |
| 5BTP | A | ZTP riboswitch |
| 5CCB | N | tRNA3Lys |
| 5CZZ | B | RNA |
| 5D8H | A | 23S ribosomal RNA |
| 5DDP | A | L-glutamine riboswitch |
| 5DEA | A | sc1 |
| 5DM6 | Y | 5S rRNA |
| 5DQK | A | Hammerhead ribozyme |
| 5E6M | E | tRNA-GLY |
| 5EL6 | 3K | tRNA-LYS |
| 5F9R | A | sgRNA |
| 5FLX | z | HCV-IRES |
| 5G2X | A | Group IIA Intron |
| 5GAN | V | U4 snRNA |
| 5J01 | A | Group IIC intron lariat |
| 5J8B | x | P-site tRNA |
| 5JUP | E | IRES |
| 5K7D | A | Pistol ribozyme |
| 5KK5 | B | crRNA |
| 5KPY | A | 5-hydroxytryptophan RNA aptamer |
| 5LYU | A | 7SK RNA |
| 5LZD | y | Sec-tRNASec |
| 5LZS | 2 | tRNA |
| 5M0H | A | ASH1 E3 (42 nt-TL/TLR) |
| 5ML7 | A | 23S ribosomal RNA |
| 5MMI | B | 5S rRNA |
| 5NWQ | A | Guanidine III riboswitch |
| 5O2R | x | P-site tRNA-ILE |
| 5O60 | B | 5S rRNA |
| 5OB3 | A | RNA aptamer |

**Table 3.** (Continued.)

| PDB ID | Chain ID | Description |
|---|---|---|
| 5ON2 | E | tRNA-LEU |
| 5OQL | 2 | U3 snoRNA |
| 5T2A | E | srRNA1 |
| 5T5A | A | Twister Sister (TS) Ribozyme |
| 5T5H | D | 5S rRNA |
| 5T83 | A | Guanidine-I riboswitch |
| 5TF6 | B | U6 snRNA |
| 5TPY | A | Exonuclease resistant RNA |
| 5U33 | B | sgRNA |
| 5U3G | B | ykkC riboswitch |
| 5U4J | a | 16S rRNA |
| 5UD5 | C | tRNA-PYL |
| 5UQ8 | x | mRNA |
| 5V3F | A | Fluorogenic RNA Mango |
| 5V3I | A | VS Ribozyme RNA |
| 5VOE | A | Aptamer 11F7t |
| 5VPP | QV | P-site tRNA SufA6 |
| 5VT0 | R | 6S RNA derivative |
| 5WLC | L0 | 5’ external transcribed spacer (5’ ETS) |
| 5WLC | L2 | U3 snoRNA |
| 5WT3 | C | tRNA |
| 5WTI | B | RNA |
| 5WWR | C | tRNA |
| 5X2H | B | sgRNA |
| 5X6B | P | tRNA-CYS |
| 5XTM | B | RNA fragment containing a K-turn motif |
| 5XWY | B | crRNA for Cas13a |
| 5XXB | 3 | 5S RNA |
| 5XXB | 4 | 5.8S RNA |
| 5XY3 | 3 | 5S rRNA |
| 5XY3 | 4 | 5.8S rRNA |
| 5Y58 | X | TLC1 |
| 5Y7M | B | RNA fragments containing a K-turn motif |
| 5Y85 | B | Four-way junctional Twister-Sister ribozyme |
| 5ZLU | X | 5S rRNA |
| 5ZTM | C | Non-coding mRNA sequence roX2 |
| 5ZWN | P | U1 snRNA |
| 6AAY | B | crRNA for Cas13b |
| 6AGB | A | Ribonuclease P RNA |
| 6AH3 | T | pre-tRNA |
| 6AHD | I | U4snRNA |
| 6AHD | B | U5snRNA |
| 6AHU | A | H1 RNA (the RNA component of RNase P) |
| 6AZ1 | 2 | tRNA-PHE |
| 6AZ3 | 4 | rRNA delta |
| 6AZ3 | 5 | rRNA epsilon |
| 6AZ3 | 6 | rRNA zeta |
| 6AZ3 | 7 | rRNA 5.8S |
| 6B14 | R | RNA aptamer |
| 6C0F | 6 | ITS2 |
| 6C65 | A | Mango-II-A22U Fluorescent Aptamer |
| 6CAE | 1w | A-site and E-site tRNAs |
| 6CF2 | G | RNA aptamer |
| 6CHR | A | Group IIB intron lariat |
| 6CK5 | A | PRPP riboswitch |
| 6CU1 | A | YrlA effector-binding module |
| 6D12 | C | 7SK RNA stem-loop 4 |
| 6D3P | A | Exoribonuclease-resistant RNA from Sweet clover necrotic mosaic virus |
| 6D90 | 4 | CrPV-IRES |
| 6DB8 | R | DIR2s RNA aptamer |
| 6DCB | B | 7SK RNA stem-loop 1 proximal |
| 6DLR | A | PRPP Riboswitch |
| 6DMC | A | ppGpp Riboswitch |
| 6DTD | C | crRNA |
| 6DVK | H | Computationally designed RNA |
| 6E8S | A | iMango-III aptamer |
| 6E9E | B | crRNA |
| 6ERI | AB | 4.5S rRNA |
| 6FRK | 1 | 7SL RNA, cytoplasmic 1 (RN7SL1), SRP RNA |
| 6FT6 | 2 | 7S rRNA |

**Table 3.** (Continued.)

| PDB ID | Chain ID | Description |
|---|---|---|
| 6FZ0 | A | metY SAM V |
| 6G90 | 1 | U1 snRNA, U1 snRNA, U1 snRNA, U1 snRNA, U1 snRNA |
| 6GAW | BB | tRNA-PHE, mitochondrial |
| 6GAZ | AV | P-site fMet-tRNAMet, mitochondrial |
| 6GYV | A | Lariat-capping ribozyme |
| 6H0R | A | SRS2 fragment of Rgs4 3' UTR |
| 6HA1 | B | 5S rRNA |
| 6I0Y | V | tRNA-PRO |
| 6ID0 | H | U2snRNA |
| 6ID1 | F | U6snRNA |
| 6IP5 | zu | P-site tRNA |
| 6IV8 | B | crRNA for Cas13d |
| 6J6G | D | U5 snRNA |
| 6J6G | E | U6 snRNA |
| 6JDV | B | sgRNA |
| 6JOO | B | Guide RNA |
| 6JQ5 | A | Hatchet Ribozyme |
| 6JXM | B | RNA |
| 6LAX | B | SAM-VI riboswitch |
| 6LXD | D | Pri-miRNA |
| 6MJ0 | A | Turnip yellow mosaic virus 3'UTR |
| 6MWN | A | Hepatitis A virus IRES domain V |
| 6N2V | A | Mn riboswitch optimized construct |
| 6N5Q | A | pir-miRNA-378a apical loop and one-base-pair fused to YdaO riboswitch |
| 6N7R | R | U1 snRNA |
| 6NY2 | B | RNA |
| 6OL3 | C | Adenovirus Virus-Associated (VA) RNA I apical and central domains |
| 6OLE | D | 5.8S rRNA |
| 6P2H | A | 2'-dG-II class of riboswitches |
| 6P5N | 1 | IAPV-IRES |
| 6PMO | A | T-box riboswitch discriminator |
| 6Q95 | 4 | transfer-messenger RNA (tmRNA) |
| 6Q97 | 7 | tRNA-VAL |
| 6Q9A | 4 | tmRNA |
| 6Q9A | 7 | P-site tRNA |
| 6QDW | v | tRNA-GLY |
| 6QN3 | A | Glutamine II Riboswitch |
| 6QX9 | 1 | U1 snRNA |
| 6QZP | L7 | 5S rRNA |
| 6QZP | S6 | E site tRNA |
| 6R47 | A | Pistol ribozyme |
| 6R5Q | 2 | P-tRNA |
| 6R87 | B | tRNA-ALA |
| 6RFL | U | chr17.trna16-GlnTTG |
| 6RJA | D | sgRNA |
| 6RM3 | L70 | 5S rRNA |
| 6S0X | B | 5S rRNA |
| 6SWE | 2 | 16S rRNA |
| 6SY4 | C | TetR-binding aptamer K1 |
| 6T4Q | C4 | 5S rRNA |
| 6T4Q | 6 | ICG tRNA Arg (P/P) |
| 6T4Q | 7 | tRNA (E/E) |
| 6T7T | 6 | tRNA |
| 6TB3 | n | tRNA |
| 6TBV | PTR1 | P-site tRNA-ARG |
| 6U8D | A | JIIIabc RNA |
| 6UES | A | Apo SAM-IV Riboswitch |
| 6UFH | A | ileS T-box |
| 6UFJ | C | Pistol ribozyme product |
| 6UFM | A | RNA |
| 6UGG | A | tRNA-ASP |
| 6V3A | v | tRNA-MET |
| 6V3A | B | 5s rRNA |
| 6V5B | D | Pri-miR-16-2 |
| 6W2S | 0 | CrPV 5'-UTR IRES |
| 6XYW | 3 | Plant mitochondrial rRNA |

**Table 4.** Detailed list of 13 RNA PDB IDs for training-like test set.

| PDB ID | Chain ID | Description |
|---|---|---|
| 7WIF | V | THF-II riboswitch |
| 8K30 | B | medaka mascRNA U23G |
| 8SA3 | B | Adenosylcobalamin-bound riboswitch dimer, form 2 |
| 8TJX | N | Tetrahymena Ribozyme |
| 8ZMI | C | BMV TLS-TyrRS-ATP(Pre-1a state) |
| 9BZ1 | A | Clostridium beijerinckii ZTP riboswitch |
| 9DE5 | D | HIV TAR RNA |
| 9G4Q | A | salivarius-1 RNA motif |
| 9NZT | E | Guide RNA |
| 9OY2 | A | RNase P ribozyme tetraloop mutant |
| 9QTJ | 1 | Oceanobacillus iheyensis group II intron domain |
| 9U5Q | R | cloverleafRNA-conformation1 |
| 9V4V | A | Guanine-II riboswitch |

**Table 5.** Detailed list of 15 RNA PDB IDs for novel test set.

| PDB ID | Chain ID | Description |
|---|---|---|
| 6XJZ | A | self-alkylating ribozyme - apo form |
| 7V9E | A | methyl transferase ribozyme |
| 8BF8 | B | reRNA |
| 8HBA | B | NAD-II riboswitch |
| 8HUJ | C | IRES RNA (J-K-St) |
| 8UYG | A | BtCoV-HKU5 5' proximal stem-loop 5, conformation 2 |
| 8VCI | A | SARS-CoV-2 Frameshift Stimulatory Element with Upstream Multibranch Loop |
| 8XZW | A | THF-II riboswitch |
| 9C2K | C | HIV-1 Rev Response Element Stem-Loop II |
| 9CFN | A | exoribonuclease-resistant RNA |
| 9DIB | A | Rous sarcoma virus frameshifting pseudoknot RNA |
| 9E73 | A | RaiA RNA |
| 9E9O | A | SARS-CoV-2 SL5 |
| 9LEC | J | Sag-18RS21 Golld RNA |
| 9MCW | A | OLE RNA dimer |
